# Hierarchical non-negative matrix factorization using clinical information for microbial communities

**DOI:** 10.1101/690123

**Authors:** Ko Abe, Masaaki Hirayama, Kinji Ohno, Teppei Shimamura

## Abstract

**Background:** The human microbiome forms very complex communities that consist of hundreds to thousands of different microorganisms that not only affect the host, but also participate in disease processes. Several state-of-the-art methods have been proposed for learning the structure of microbial communities and to investigate the relationship between microorganisms and host environmental factors. However, these methods were mainly designed to model and analyze single microbial communities that do not interact with or depend on other communities. Such methods therefore cannot comprehend the properties between interdependent systems in communities that affect host behavior and disease processes.

**Results:** We introduce a novel hierarchical Bayesian framework, called BALSAMICO (BAyesian Latent Semantic Analysis of MIcrobial COmmunities), which uses microbial metagenome data to discover the underlying microbial community structures and the associations between microbiota and their environmental factors. BALSAMICO models mixtures of communities in the framework of nonnegative matrix factorization, taking into account environmental factors. This method first proposes an efficient procedure for estimating parameters. A simulation then evaluates the accuracy of the estimated parameters. Finally, the method is used to analyze clinical data. In this analysis, we successfully detected bacteria related to colorectal cancer. These results show that the method not only accurately estimates the parameters needed to analyze the connections between communities of microbiota and their environments, but also allows for the effective detection of these communities in real-world circumstances.

## Background

Microbiota in the human gut form complex communities that consist of hundreds to thousands of dierent microorganisms that aect various important functions such as the maturation of the immune system, physiology [1], metabolism [2], and nutrient circulation [3]. Species in a community survive by interacting with each other and can concurrently belong to multiple communities [4]. Moreover, the composition of bacterial species can change over time. In some cases, a single species or strain significantly affects the state of the community, making it a causative agent for disease. For example, *Helicobacter pylori* is a pathogen that induces peptic disease [5]. However, problems are not always rooted in an individual species or strain. In many cases it is the dierences in dierent types of microbial communities, i.e. their composition ratios, that affect the overall structure of the gut microbiota. These overall structures relate to various features of interest— for example, the ecosystem process [6], the severity of the disease [7], or the impact of dietary intervention [8]. Therefore, finding co-occurrence relationships between species and revealing the community structure of microorganisms is crucial to understanding the principles and mechanisms of microbiota-associated health and disease relationships and interactions between the host and microbe.

Thanks to modern technology, revealing these community structures is becoming easier. Advances in high-throughput sequencing technologies such as shotgun metagenomics have made it possible to investigate the relationship among microorganisms within the whole gut ecosystem and to observe the interaction between microbiota and their host environments. Many microbiome projects, including the Human Microbiome Project (HMP) [9] and the Metagenomics and the Human Intestinal Tract (MetaHIT) project [10], have generated considerable data regarding human microbiota by studying microbial diversity in different environments. The data consists of either marker-gene data (the abundance of operational taxonomic units; OTUs) or functional metagenomic data (the abundance of reaction-coding enzymes). Although collecting such data is no longer methodologically difficult, analysis remains challenging. Even with limited samples, the data always consists of hundreds or even thousands of variables (OTUs or enzymes). In addition, there are many rare species of microbiota, and these are observed only in very few samples. Thus the data is highly sparse [11]. The sparse nature of the data means that classical statistical analysis methods, which were designed for data rich situations, have limited ability to identify complex features and structures within the data. Several new methods are therefore emerging in order to properly analyze and understand microbiota.

In this study, we focus on learning the structure of microbial communities and investigating the relationship between microorganisms and their environmental factors using metagenomic data. Currently, there are several methods that seek to clarify this relationship. One is probabilistic modeling of metagenomic data, which often provides a powerful framework for the problem. For example, [12] proposed BioMiCo, a two-level hierarchical Bayes model of a mixture of multidimensional distributions constrained by Dirichlt priors to identify each OTU cluster, called an assemblage, and to estimate the mixing ratio of the assemblages within a sample. Another popular method for learning community structure is non-negative matrix factorization (NMF) [13, 14]. Cai *et al*. [15] proposed a supervised version of NMF to identify communities representing the connection between the sample microbial composition and OTUs and to infer systematic dierences between dierent types of communities. These methods are useful in a variety of circumstances, but they also possess limitations.

When it comes to learning the structure of microbial communities related to environmental features of interest, the limitations of the current approaches become clear. Although BioMiCo can learn how microbes contribute to an underlying community structure that is related to a known feature of each sample, it fails when the microbes are composed of a mixture of communities that interact with each other. In such cases, another method must be applied. Supervised NMF is one option, as it can be used to extract communities that are characterized by a co-occurrence relationship. However, in this framework, the analyst must explicitly specify the communities to which the bacteria belong [15]. This process depends on the knowledge of the analyst, so in cases of limited information about the communities the method cannot be used. To our knowledge, no framework currently exists that adequately details the interaction between a mixture of microbial communities and multiple environmental factors. A new framework is needed to address this problem.

To remedy this situation, we propose a novel approach, called BALSAMICO (BAyesian Latent Semantic Analysis of MIcrobial COmmunities). The contributions of our research are as follows:

- BALSAMICO uses the OTU abundances and the host environmental factors as input to provide a path to interpret microbial communities and their environmental factors. In BALSAMICO, the data matrix of a microbiome is approximated to the product of two matrices. One matrix is represented by a mixing ratio of microbial communities, and the other matrix is represented by the abundance of bacteria in the communities. BALSAMICO decomposes the mixing ratio into the observed environmental factors and their coefficients in order to identify the influence of the environmental factors.
- Not only is this decomposition a part of ordinary NMF, but it improves upon ordinary NMF by displaying a hierarchical structure. One clear advantage of the Bayesian hierarchical model is to introduce stochastic fluctuations at all levels. This makes it possible to smoothly handle missing data and to easily give credible intervals.
- Unlike supervised NMF, BALSAMICO does not require prior knowledge regarding the communities to which the bacteria belong. BALSAMICO can estimate an unknown community structure without explicitly using predetermined community information. Furthermore, the parameters of unknown community structures can be estimated automatically through Bayesian learning.
- While the computation cost of other methods, which use Gibbs sampling, is high, we provide an efficient learning procedure for BALSAMICO by using a variational Bayesian inference and Laplace approximation to reduce computational cost. The software package that implements BALSAMICO in the R environment is available from GitHub (https://github.com/abikoushi/BALSAMICO).

The structure of this paper will proceed as follows: The “Methods” section describes our model and the procedure for parameter estimation. The “Results” section contains an evaluation of the accuracy of the estimator using synthetic data. Additionally, BALSAMICO is applied to clinical metagenomic data to detect bacterial communities related to colorectal cancer (CRC). Through this content, both the usefulness and accuracy of BALSAMICO are confirmed.

## Methods

Calculations for this method are based on the assumption that the microbiome consists of several communities. BALSAMICO extracts the communities from the data, using NMF. Suppose that we observe a nonnegative integer matrix ***Y*** = (*y*_*n,k*_) (*n* = 1,…, *N, k* = 1,…, *K*), where *y*_*n,k*_ is the microbial abundance of *k*-th taxon in the *n*-th sample. Our goal is to seek a positive *N* × *L* matrix ***W*** and an *L* × *K* matrix ***H***, such that

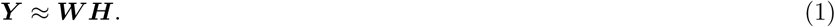

The (*n, l*)-element *w*_*n,l*_ of matrix ***W*** can be interpreted as contributing to community *l* of sample *n*. The (*l, k*)-element *h*_*l,k*_ of matrix ***H*** can be interpreted as the relative abundance of the *k*-th taxon given community *l*. We thus refer to ***W*** as the *contribution matrix* and to ***H*** as the *excitation matrix*.

In addition, if covariate ***X*** = (*x*_*n,d*_) (*d* = 1,…, *D*) is observed (e.g. whether or not the *n*-th sample has a certain disease), our aim is to seek how ***W*** changes when ***X*** is given. For this, BALSAMICO seeks the *D* × *L* matrix ***V***, such that

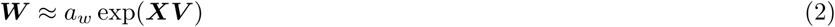

where exp(·) is an element-wise exponential function. As shown in Figure 1, BALSAMICO approximates matrix ***Y*** using the product of low-rank matrices.

**Figure 1.**
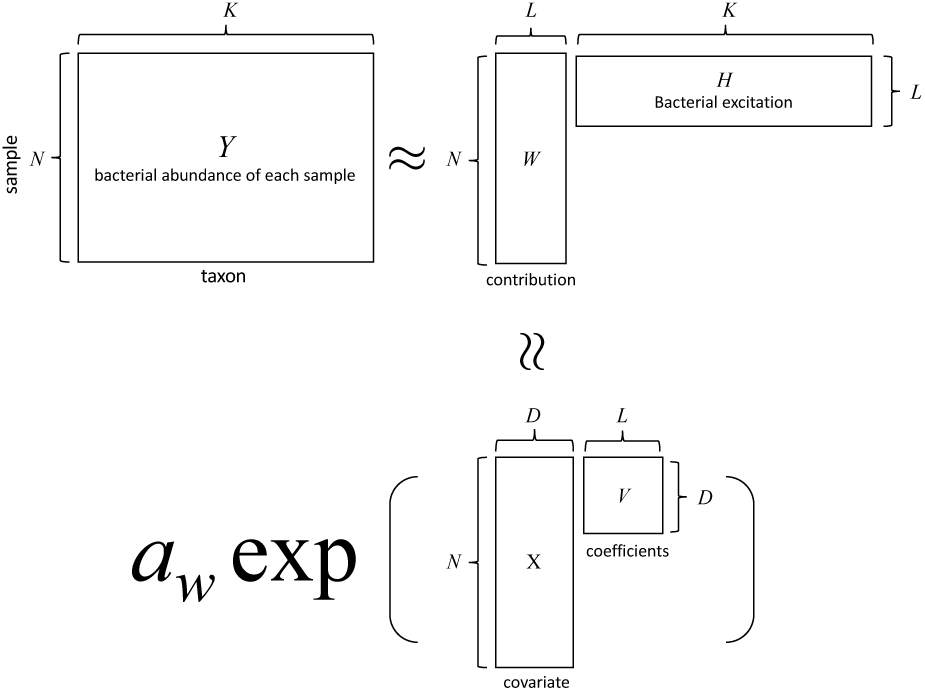
Conceptual diagram of matrix factorization in BALSAMICO.

In brief, we consider the following hierarchical model:

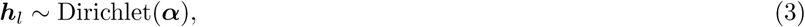

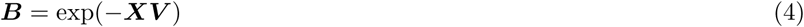

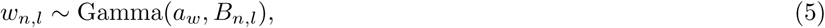

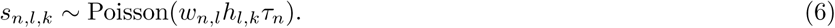

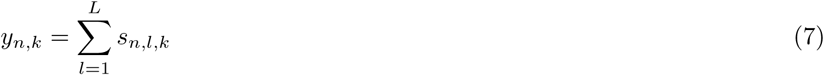

where *B*_*n,l*_ is the (*n, l*)-element of matrix ***B***, *τ*_*n*_ is an oset term, ***V*** is a *D*×*L* matrix, and ***S*** = {*s*_*n,l,k*_} are latent variables. The variable ***S*** is introduced for inference to make the calculations more smooth. In this study, we set 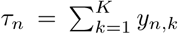. The total read count *τ*_*n*_ is dependent on the setting of the DNA sequencer, so it is not a reflection of an abundance of bacteria. The offset term then adjusts the setting-based effect on the read counts to accurately estimate ***W***. The (*d, l*)-element *v*_*d,l*_ of matrix ***V*** can be interpreted as contributing to the community *l* of the *d*-th covariate. This Poisson observation model is frequently used in Bayesian NMF [16]. Gamma and Dirichlet prior distribution are the conjugate priors.

Figure 2 shows a plate diagram of the data generating process. BALSAMICO estimates parameters ***W***, ***H***, *a*_*w*_, and ***V***, using variational inference [17]. More details for this parameter estimation procedure are listed in the supplemental document. After estimating the parameters it is possible to move on to analyzing real data, but first the accuracy of the estimation should be confirmed.

**Figure 2.**
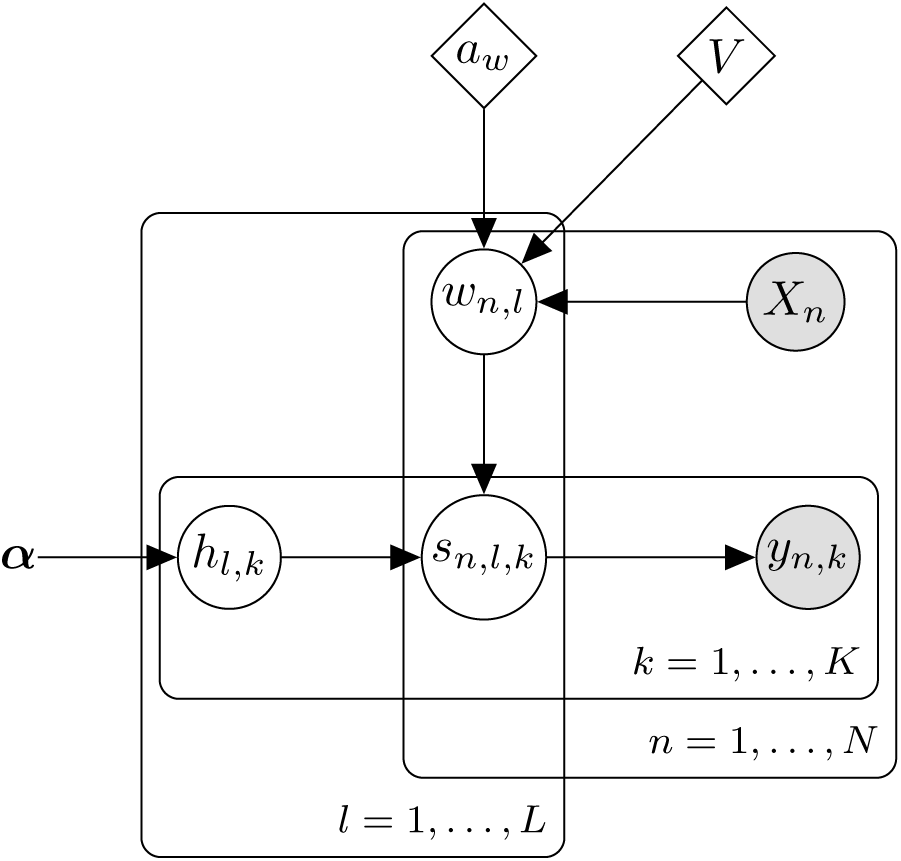
Plate diagram of the data generating process in BALSAMICO. The white nodes indicate latent variables and the gray nodes indicate observed variables. The parameters represented by diamonds are estimated by Laplace approximation.

## Results

### Simulation Study

Starting with the BALSAMICO estimated parameters detailed in “Methods,” we can now evaluate these parameters for accuracy before moving on to an analysis of real-world data. The following simulation experiments evaluate the bias, the standard error (SE), and the coverage probability (CP) of the estimators. The bias of 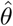 is defined by the difference between the true value and the estimated value 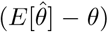. The coverage probability is the proportion at which the 95% credible interval contains the true value. The synthetic data was naturally produced via the data generating process given by Eqs. 3–7.

We estimated the parameters in 10,000 replicates of the experiment. We set *X* = (**1**, ***x***_1_, ***x***_2_), where **1** is a vector of ones. The variables ***x***_1_ and ***x***_2_ are sampled from the standard normal distribution and the Bernoulli distribution with a probability of 0.5. When generating the synthetic data, we set *N* = 100, *K* = 100, *L* = 3, *τ*_*n*_ = 10, 000, and *α*_*k*_ = 1 for all *k*. We also set *α*_*k*_ = 1 for all *k* when estimating parameters, which is equivalent to a non-informative prior distribution. To avoid the problem of label switching [18], the estimated parameters are rearranged as *v*_21_ ≤ *v*_22_ ≤ *v*_2_3.

The gamma distribution changes considerably when the shape parameter *a*_*W*_ is smaller than 1, which leads to a heavier tail than an exponential distribution. Consequently, we conducted two patterns of the simulation. Table 1 shows these results. The first half of the table shows the case of a heavy tail.

**Table 1.** Bias, se, and CP of the estimates

|  | true value | bias | se | CP |
| --- | --- | --- | --- | --- |
| $a_w$ | 0.5 | -0.01 | 0.10 | |
| $v_{11}$ | 1.00 | 0.00 | 0.30 | 0.86 |
| $v_{12}$ | -0.50 | -0.00 | 0.15 | 0.95 |
| $v_{13}$ | 0.50 | 0.00 | 0.30 | 0.94 |
| $v_{21}$ | 1.00 | 0.01 | 0.30 | 0.86 |
| $v_{22}$ | 0.00 | -0.00 | 0.15 | 0.95 |
| $v_{23}$ | 0.00 | 0.00 | 0.30 | 0.94 |
| $v_{31}$ | 1.00 | 0.01 | 0.30 | 0.86 |
| $v_{32}$ | 0.50 | 0.00 | 0.15 | 0.95 |
| $v_{33}$ | -0.50 | 0.01 | 0.29 | 0.95 |
| $a_w$ | 2.00 | 0.06 | 0.17 | |
| $v_{11}$ | 1.00 | -0.04 | 0.13 | 0.93 |
| $v_{12}$ | -0.50 | -0.00 | 0.07 | 0.94 |
| $v_{13}$ | 0.50 | 0.00 | 0.15 | 0.94 |
| $v_{21}$ | 1.00 | -0.04 | 0.13 | 0.92 |
| $v_{22}$ | 0.00 | 0.00 | 0.07 | 0.94 |
| $v_{23}$ | 0.00 | 0.00 | 0.15 | 0.94 |
| $v_{31}$ | 1.00 | -0.03 | 0.13 | 0.94 |
| $v_{32}$ | 0.50 | -0.00 | 0.07 | 0.94 |
| $v_{33}$ | -0.50 | 0.01 | 0.15 | 0.95 |

When the shape parameter *a*_*W*_ is set to 0.5, the credible intervals of *v*_*i*1_ (*i* = 1, 2, 3) have under-coverage. However, this was only observed in intercept terms. In most cases, the CP was an almost nominal value. This result indicates that there is no inconsistency when interpreting the estimated coefficients.

Moreover, the parameters were estimated with small biases. By this we know that the proposed method produces reasonable estimates. This being confirmed, it is now possible to apply the proposed method to real data to assess how well it conforms to current studies.

### Results on real data

This section tests the usefulness of our results by investigating the identification of gut dysbiosis associated with the development of CRC. Zeller *et ql*. [19] studied gut metagenomes extracted from 199 persons: 91 CRC patients, 42 adenoma patients, and 66 controls. The data is available in the R package “curatedMetage-nomicData” (https://github.com/waldronlab/curatedMetagenomicData). This analysis uses the abundance of genus-level taxa.

We set *α*_*k*_ = 1 and use the disease label, gender, and age as covariates. The age variable is scaled by dividing by 100. The number of communities *L* = 7 was selected using leave-one-out cross-validation (Figure 3).

**Figure 3.**
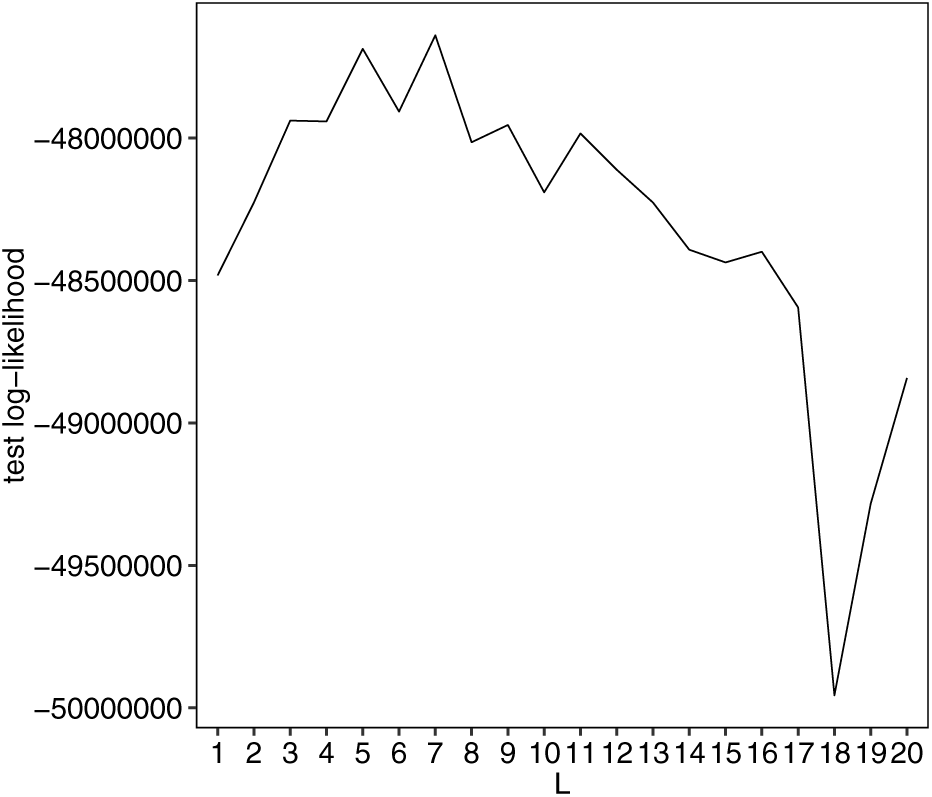
Mean of test log-likelihood evaluated by leave-one-out cross-validation. The *x*-axis corresponds to the number of communities *L*.

Figure 4 shows the estimated ***WH*** and normalized abundance 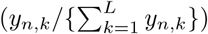. The observed data matrix is approximated by ***WH***.

**Figure 4.**
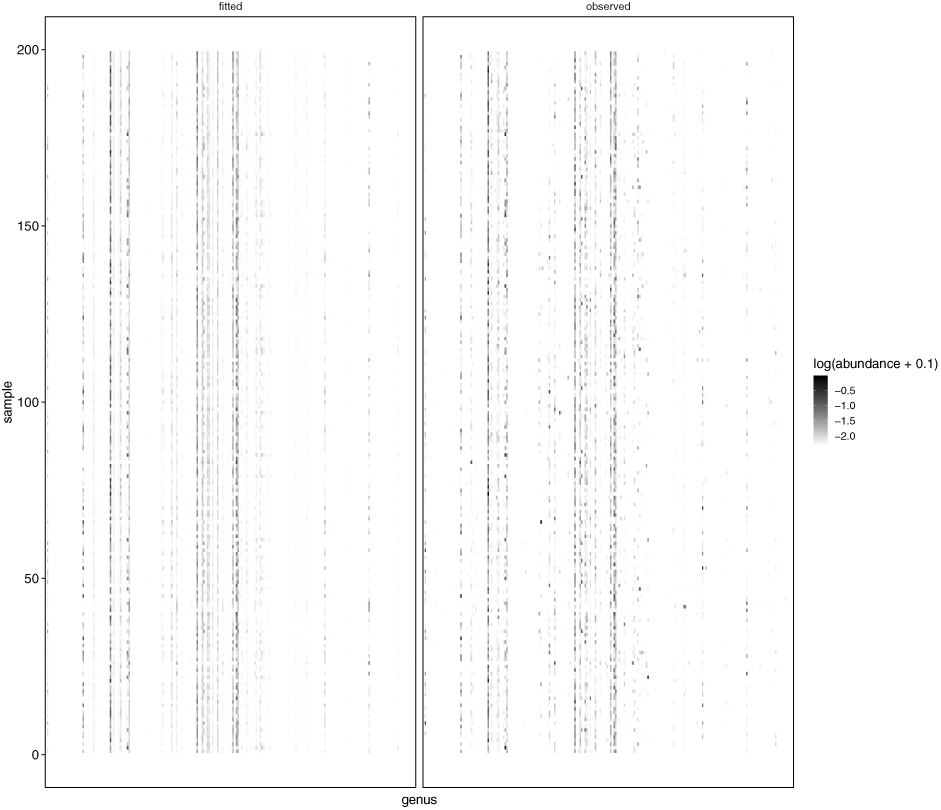
Comparison between *W H* (fitted) and normalized abundance (observed). For visibility, we use a logarithmic scale. The *x*-axis corresponds to the genera, and the *y*-axis corresponds to the samples.

**Figure 5.**
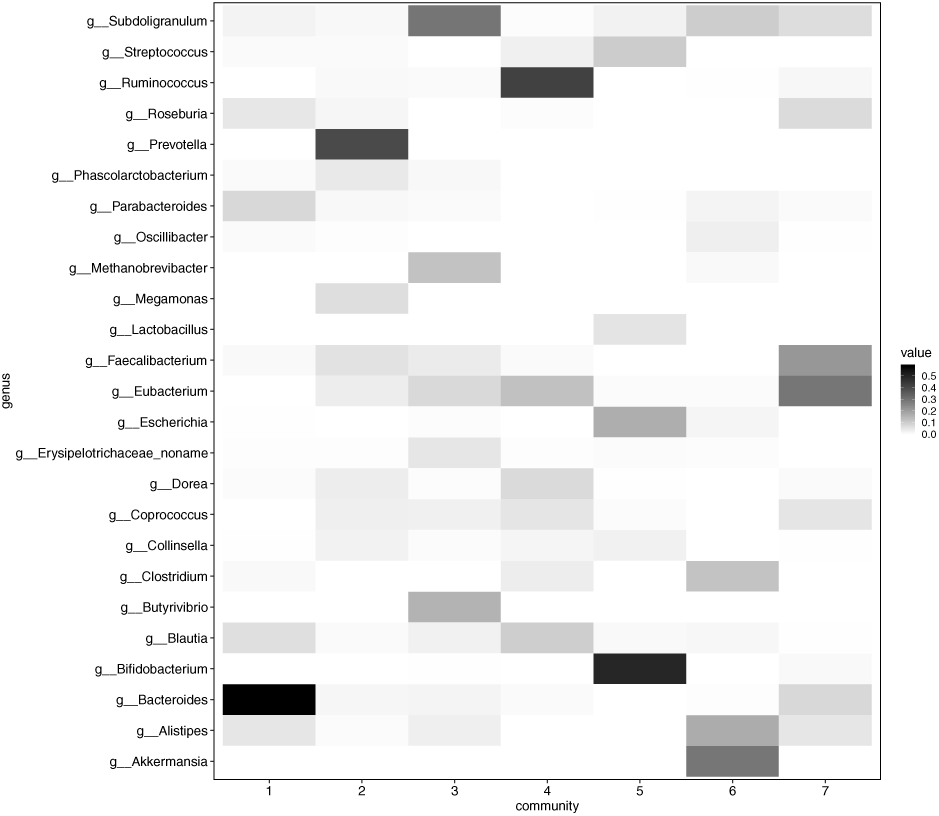
Estimated excitation matrix *H*. The *x*-axis corresponds to the community, and the *y*-axis corresponds to the genus. The black parts indicate high abundance, and the white parts indicate zero.

Figure 6 shows estimates of coefficient ***V***. First, we can see that the human microbiome is not dependent on gender as the absolute value of coefficients for gender is small, and their credible intervals contain zero. Focusing on CRC, we can see that the credible intervals of the coefficient for community 6 do not contain zeros. Moreover the value of coefficients for community 6 increases as adenoma progresses to CRC. Community 6 is thus strongly suspected of being associated with the disease. Figure 7 shows estimates of *W*_*n*,6_. We observed individual differences, but, overall, CRC patients have large community 6, which confirms this suspicion.

**Figure 6.**
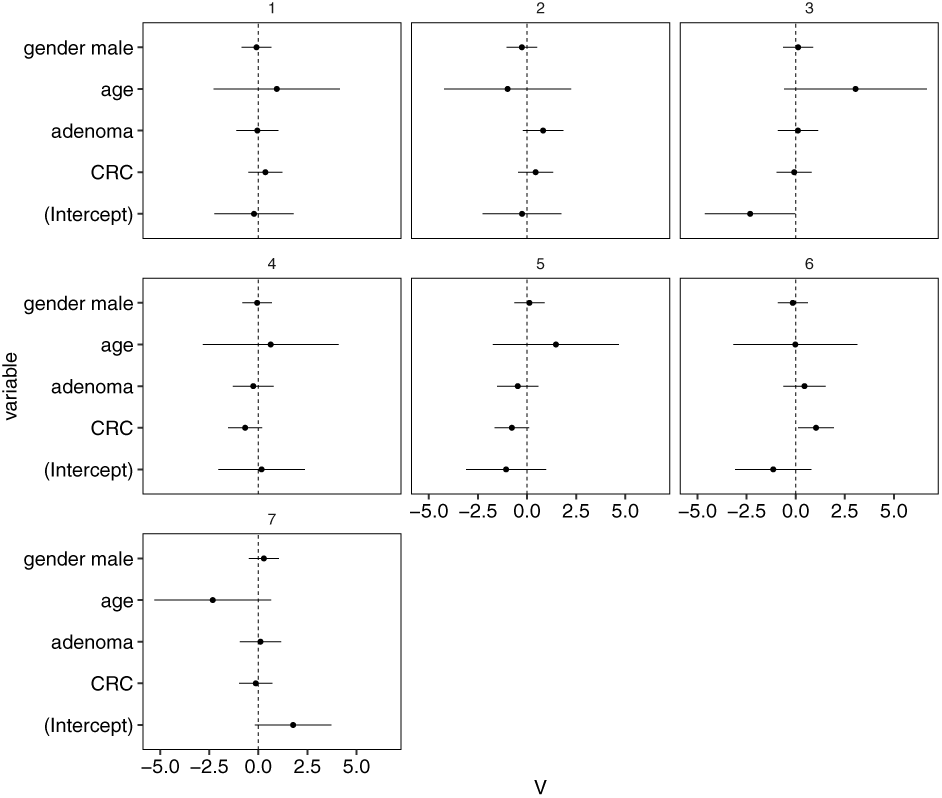
Estimated coefficients *V* for the disease label. The each panel corresponds to community, the *x*-axis corresponds to the value of coefficients and the *y*-axis corresponds to the variable name. The bars indicate 95%-credible intervals.

**Figure 7.**
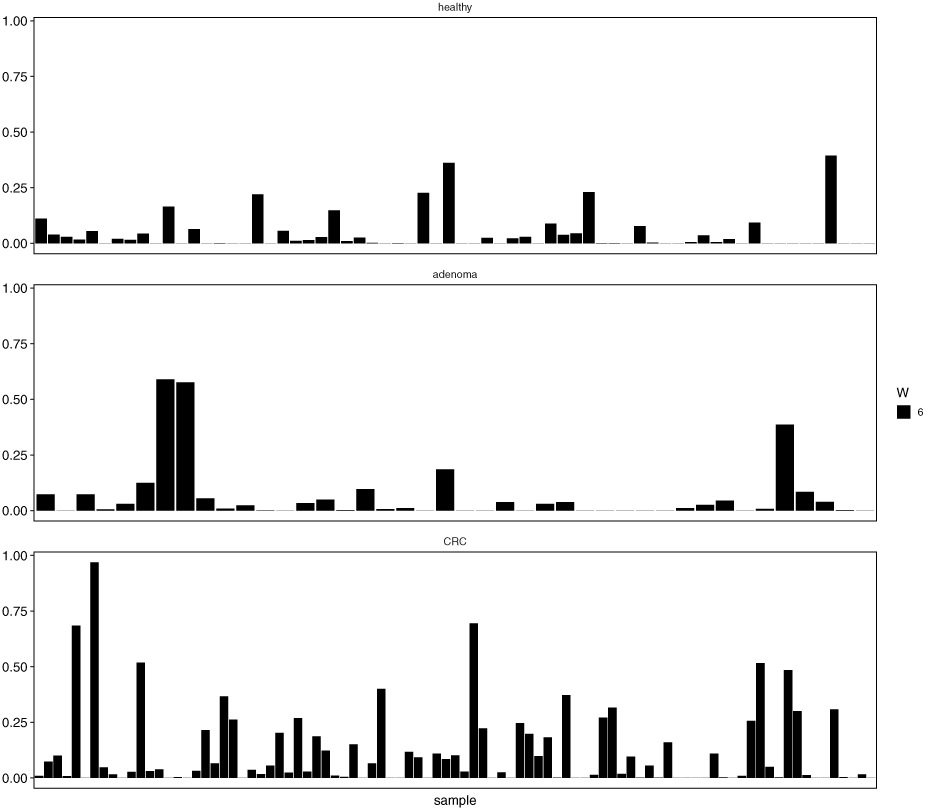
Estimated contribution matrix *w*_*n*,6_ of each sample *n*. The *x*-axis corresponds to the samples, and the *y*-axis corresponds to the value of *w*_*n*,6_.

Figure 5 shows the top five estimates of *h*_*l,k*_ in each community *l*. Arumugam *et al*. reports that the human gut microbiome can be classified into several types, called enterotypes. Arumugam *et al*. [20] shows that an enterotype is characterized by the differences in the abundance of *Bacteroides, Prevotella*, and *Ruminococcus*. Communities 1, 2, and 4 are characterized by an abundance of *Bacteroides, Prevotella*, and *Ruminococcus* respectively (Figure 5). Communities 1, 2, and 4 thus correspond to enterotypes. Community 6, which is suspected of being related to CRC, is characterized by abundant *Akkermansia*. This is markedly different from the other communities and deserves further examination.

To detect the bacteria that exists exclusively in community 6, we use following amount:

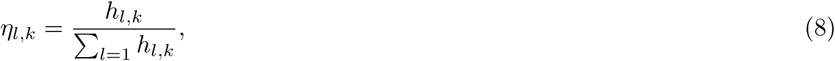

where *η*_*l,k*_ is the probability that a certain taxon *k* belongs to community *l*.

The bacteria belonging to community 6 are suspected of being associated with CRC. Table 2 shows estimates of *η*_6,*k*_ greater than 0.95. This result indicates that these bacteria are related to CRC. These bacteria that characterize community 6 are *Akkermansia, Desulfotomaculum, Mucispirillum, Methanobacterium, Hahellaceae, Nakaseomyces, Fretibacterium, Alphabaculovirus, Synergistes*, and *Enhydrobacte*. The connection between these bacteria and CRC is further supported by current studies.

**Table 2.**
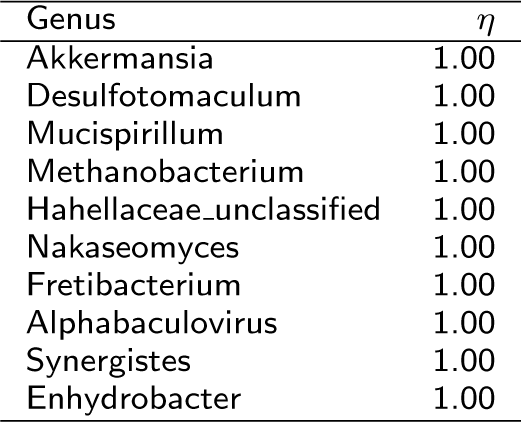
Estimates of *η*_6,*k*_ greater than 0.95

| Genus | $\eta$ |
| --- | --- |
| Akkermansia | 1.00 |
| Desulfotomaculum | 1.00 |
| Mucispirillum | 1.00 |
| Methanobacterium | 1.00 |
| Hahellaceae_unclassified | 1.00 |
| Nakaseomyces | 1.00 |
| Fretibacterium | 1.00 |
| Alphabaculovirus | 1.00 |
| Synergistes | 1.00 |
| Enhydrobacter | 1.00 |

- *Akkermansia*: Weir *et al*. [21] reports that mucin-degrading bacteria, *Akkermansia muciniphila*, was present in a significantly greater proportion in the feces of colon cancer patients. This is consistent with our result.
- *Desulfotomaculum*: *Desulfotomaculum* belongs to sulfate-reducing bacteria, which obtains energy by oxidizing organic compounds or molecular hydrogen while reducing sulfate to hydrogen sulfide. Hydrogen sulfide is toxic to intestinal epithelium cells and causes DNA damage in human cells [22].
- *Mucispirillum*: Similar to *Akkermansia, Mucispirillum* is a mucus-resident bacteria and may coexist with *Akkermansia*. If so, these bacteria are distributed in mucus layer that covers the mucous membrane of the intestine. [23].
- *Methanobacterium*: Patients with CRC contain a higher proportion of breath methane excreters than the control group [24]. *Methanobacterium* is a methanogenic bacterium.
- *Enhydrobacter*: Xu & Jiang [25] uses linear discriminative analysis to biomarker discovery. The result suggests that *Enhydrobacter* can be a biomarker for CRC.

The information found in the above studies strongly supports the results returned by applying our method to real data. This suggests that BALSAMICO is able to successfully and accurately analyze communities of bacteria and their environmental interactions.

The information found in the above studies strongly supports the results returned by applying our method to real data. This suggests that BALSAMICO is able to successfully and accurately analyze communities of bacteria and their environmental interactions.

## Conclusion

We proposed a novel hierarchical Bayesian model to discover the underlying microbial community structures and the associations between microbiota and their environmental factors based on microbial metagenomic data. One of the most important features of our model is to decompose the contribution matrix into observed environmental factors and their coefficients. The parameters for this model were estimated using variational Bayesian inference, as described in “Methods”. In terms of computation, this parameter-estimation procedure offers two advantages over existing methods. First, in an algorithm that uses Gibbs sampling, the computational cost is large due to the large number of samples required. By contrast, our procedure involves a matrix operation that substitutes for this requirement, helping to reduce computational cost. Second, our procedure involves hyper-parameter tuning. The parameters of the gamma prior distribution are estimated from the data. The parameters of the Dirichlet prior distribution can be non-informative, and the number of communities *L* can be selected by cross-validation.

The results of our simulations suggest that the estimators of the effects of environmental factors ***V*** are consistent. Generally, other NMF methods lack consistency because they may not have a unique solution [15]. Indeed, the consistency of our method increases the reproducibility of the analysis. Moreover, the credible intervals of coefficient ***V*** are easily computed and help to identify notable bacteria.

From the perspective of data analysis, BALSAMICO has useful properties. Using the Dirichlet prior distribution, the excitation matrix ***H*** is easily interpreted as a relative abundance of species in communities. As shown in Figure 5, *h*_*l,k*_ obtains a value that is often close to zero. This property thus expresses data sparsity. Furthermore, the Poisson observation model may be applicable to counting other data (for example, gene expression data). The hierarchical structure of our model allows it to capture (*i*) dependencies between environmental factors and the community structure (represented by coefficient ***V***), and (*ii*) the individual differences in microbial composition (represented by the contribution matrix ***W***). Thus, BALSAMICO can be used to find latent relationships between bacteria. As discussed in “Results,” BALSAMICO’s findings from real data are supported by previous studies. This demonstrates that BALSAMICO is effective at knowledge discovery.

This research has possibility for expansion and may provide positive contributions to future studies. In many situations, microbiome data is obtained as time a series which repeats measurements for each sample. To handle the time series data, our model could be expanded so the contribution matrix ***W*** is extended from a matrix to a tensor. This facilitates the analysis of time-varying bacteria compositions during the progression of a disease. Furthermore, although this research was limited to the study of the gut microbiome in connection to CRC, BALSAMICO will prove useful to other studies seeking to find relationships between various microbiomes and environmental factors. This will allow for a better understanding of the cause of disease and how disease is impacted by the microbiome environment.

## Ethics approval and consent to participate

Not applicable.

## Consent for publication

Not applicable.

## Availability of data and material

BALSAMICO is implemented with R and is available from GitHub (https://github.com/abikoushi/BALSAMICO). The data is available in the R package “curatedMetagenomicData” (https://github.com/waldronlab/curatedMetagenomicData).

## Author’s contributions

KA and TS designed the proposed algorithm. KO and MH designed the experiments. All authors have read and approved the final manuscript.

## Acknowledgements

Not applicable.

## Additional Files

Additional file 1 — Supplemental methods

Details of variational inference.

