## Supplemental Methods for "Hierarchical non-negative matrix factorization using clinical information for microbial communities"

#### 1. Probabilistic distributions

In this article, the probability function of a Poisson distribution is defined by

$$p(k|\lambda) = e^{-\lambda}\frac{\lambda^k}{k!}. \tag{1}$$

The density function of a gamma distribution is defined by

$$p(x|\alpha, \beta) = \frac{\beta^\alpha x^{\alpha-1} e^{-\beta x}}{\Gamma(\alpha)}. \tag{2}$$

where $\Gamma(\cdot)$ is the gamma function. The density function of a Dirichlet distribution is defined by

$$p(x|\boldsymbol{\alpha}) = \frac{\prod_{k=1}^{K}\Gamma(\alpha_k)}{\Gamma(\sum_{k=1}^{K}\alpha_k)}\prod_{i=1}^{K} x_i^{\alpha_k - 1}, \tag{3}$$

where $\alpha_k$ is the $k$-th element of $\boldsymbol{\alpha}$.

#### 2. Variational Bayesian inference

The parameter-estimation procedure for BALSAMICO is based on the variational NMF [1]. However, our algorithms are not identical to those in variational NMF [1]. Ordinally, In the variational NMF, $\boldsymbol{H}$ follows a Gamma prior distribution, whereas in our study, $\boldsymbol{H}$ follows a Dirichlet prior distribution. We want to calculate the posterior distribution $q(\boldsymbol{S}, \boldsymbol{W}, \boldsymbol{H}, a_w, \boldsymbol{V})$ from our generating model given the data and hyperparameter $\boldsymbol{\alpha}$. Because this distribution is complex, however, we assume that the posterior distribution can be factorized as follows:

$$\begin{aligned} q(\boldsymbol{S}, \boldsymbol{W}, \boldsymbol{H}, \Theta) &= q(\boldsymbol{S})q(\boldsymbol{W})q(\boldsymbol{H})q(\Theta) \\ &= \left(\prod_{n=1}^{N}\prod_{k=1}^{K} q(s_{n,1:L,k})\right)\left(\prod_{n=1}^{N}\prod_{l=1}^{L} q(w_{n,l})\right)\left(\prod_{l=1}^{L}\prod_{k=1}^{K} q(h_{l,k})\right) \\ &\quad \times q(\Theta) \end{aligned} \tag{4}$$

where $\Theta = (a_w, \boldsymbol{V})$. The variational Bayes (VB) method minimizes the upper bound of the negative marginal log likelihood as follows:

$$\begin{aligned}
&-\log p(Y|\theta)\\
&= -\log \sum_{\boldsymbol{S}} \int\int\int p(\boldsymbol{Y}, \boldsymbol{S}, \boldsymbol{W}, \boldsymbol{H}, \Theta)\, d\boldsymbol{W} d\boldsymbol{H} d\Theta\\
&= -\log \sum_{\boldsymbol{S}} \int\int\int q(\boldsymbol{S}, \boldsymbol{W}, \boldsymbol{H}, \Theta)\\
&\quad \times \frac{p(\boldsymbol{Y}, \boldsymbol{S}, \boldsymbol{W}, \boldsymbol{H}, \Theta)}{q(\boldsymbol{S}, \boldsymbol{W}, \boldsymbol{H}, \Theta)}\, d\boldsymbol{W} d\boldsymbol{H} d\Theta\\
&\le -\sum_{\boldsymbol{S}} \int\int\int q(\boldsymbol{S}, \boldsymbol{W}, \boldsymbol{H}, \Theta)\\
&\quad \times \log \frac{p(\boldsymbol{Y}, \boldsymbol{S}, \boldsymbol{W}, \boldsymbol{H}, \Theta)}{q(\boldsymbol{S}, \boldsymbol{W}, \boldsymbol{H}, \Theta)}\, d\boldsymbol{W} d\boldsymbol{H} d\Theta\\
&= -\sum_{\boldsymbol{S}} \int\int q(\boldsymbol{S}) q(\boldsymbol{W}) q(\boldsymbol{H}) q(\Theta)\\
&\quad \times \log \frac{p(\boldsymbol{Y}, \boldsymbol{S}, \boldsymbol{W}, \boldsymbol{H}, \Theta)}{q(\boldsymbol{S}) q(\boldsymbol{W}) q(\boldsymbol{H}) q(\Theta)}\, d\boldsymbol{W} d\boldsymbol{H} d\Theta. \qquad (5)
\end{aligned}$$

If $q(\boldsymbol{W})$, $q(\boldsymbol{H})$, and $q(\Theta)$ are given, then Eq. (5) becomes

$$\begin{aligned}
&\mathrm{KL}(q(\boldsymbol{S}) \| \exp(\langle \log p(\boldsymbol{Y}, \boldsymbol{S}, \boldsymbol{W}, \boldsymbol{H}, \Theta) \rangle_{q(\boldsymbol{W}), q(\boldsymbol{H}), q(\Theta)})) + \text{const.}\\
&= \sum_{\boldsymbol{S}} q(\boldsymbol{S}) \log \frac{q(\boldsymbol{S})}{\exp(\langle \log p(\boldsymbol{Y}, \boldsymbol{S}, \boldsymbol{W}, \boldsymbol{H}, \Theta) \rangle_{q(\boldsymbol{W}), q(\boldsymbol{H}), q(\Theta)})}\\
&\quad + \text{const.} \qquad (6)
\end{aligned}$$

where, $\mathrm{KL}(p\|q)$ is the Kullback–Leibler divergence between $p$ and $q$, and $\langle f(x) \rangle_{p(x)}$ denotes the expectation with respect to $p(x)$. Thus, to minimize the upper bound of the negative marginal log likelihood, the update formula of the approximate distribution $q(\boldsymbol{S})$ is as follows:

$$q(\boldsymbol{S}) \propto \exp\left( \langle \log p(\boldsymbol{Y}, \boldsymbol{S}, \boldsymbol{W}, \boldsymbol{H}, \Theta | \boldsymbol{\alpha}) \rangle_{q(\boldsymbol{W}), q(\boldsymbol{H}), q(\Theta)} \right). \qquad (7)$$

Similarly, we get the update formula for $q(\boldsymbol{W})$ and $q(\boldsymbol{H})$ as follows:

$$q(\boldsymbol{W}) \propto \exp\left( \langle \log p(\boldsymbol{Y}, \boldsymbol{S}, \boldsymbol{W}, \boldsymbol{H}, \Theta | \boldsymbol{\alpha}) \rangle_{q(\boldsymbol{S}), q(\boldsymbol{H}), q(\Theta)} \right), \qquad (8)$$

$$q(\boldsymbol{H}) \propto \exp\left( \langle \log p(\boldsymbol{Y}, \boldsymbol{S}, \boldsymbol{W}, \boldsymbol{H}, \Theta | \boldsymbol{\alpha}) \rangle_{q(\boldsymbol{S}), q(\boldsymbol{W}), q(\Theta)} \right), \qquad (9)$$

$$q(\Theta) \propto \exp\left( \langle \log p(\boldsymbol{Y}, \boldsymbol{S}, \boldsymbol{W}, \boldsymbol{H}, \Theta | \boldsymbol{\alpha}) \rangle_{q(\boldsymbol{S}), q(\boldsymbol{W}), q(\boldsymbol{H})} \right). \qquad (10)$$

The update equation for $q(\boldsymbol{S})$ leads to the following:

$$\begin{aligned}
q(s_{n,1:L,k}) \propto \exp\Big( &\sum_{l=1}^{L} (s_{n,l,k} (\langle \log w_{nl} \rangle + \langle \log h_{l,k} \rangle + \log \tau_n)\\
&- \log \Gamma(s_{n,l,k} + 1)) \Big) \qquad (11)
\end{aligned}$$

where subscripts of the expectation are omitted. This is the probability function of the multinomial distribution. From the property of the multinomial distribution, the expectation of $s_{n,l,k}$ is

$$\langle s_{n,l,k} \rangle = y_{n,k} p_{n,l,k} \tag{12}$$

where

$$p_{n,l,k} = \frac{\exp(\langle \log w_{nl} \rangle) + \langle \log h_{lk} \rangle)}{\sum_{l=1}^{L} \exp(\langle \log w_{nl} \rangle) + \langle \log h_{lk} \rangle)} \tag{13}$$

.

The update equation for $q(\boldsymbol{W})$ leads to the following:

$$\begin{aligned} q(w_{n,l}) \propto \exp\Bigg( & \left( \langle a_w \rangle + \sum_{k=1}^{K} \langle s_{n,l,k} \rangle - 1 \right) \log w_{n,l} \\ & - (\langle B_{n,l} \rangle + 1) \tau_n w_{n,l} \Bigg) \end{aligned} \tag{14}$$

This is the density function of the gamma distribution.

The update equation for $q(\boldsymbol{H})$ leads to the following:

$$q(h_{l,k}) \propto \exp\left( \sum_{k=1}^{K} \left( \alpha_k + \sum_{n=1}^{N} \langle s_{n,l,k} \rangle - 1 \right) \log h_{l,k} \right). \tag{15}$$

This is the probability function of the Dirichlet distribution.

Using Laplace approximation, parameters $a_w$ and $\boldsymbol{V}$ can be estimated by maximizing Eq. (10) as follows [2]:

$$\begin{aligned} \log q(a_w, \boldsymbol{V}) = & \sum_{n=1}^{N} \sum_{l=1}^{L} (-B_{n,l} \langle w_{n,l} \rangle - a_w \log B_{n,l} \\ & + a_w \langle \log w_{n,l} \rangle - \log \Gamma(a_w)) + \text{Const.} \end{aligned} \tag{16}$$

Now, let $\rho = \log(a_w)$. We optimize Eq. (16) with a quasi-Newton method using the gradient

$$\begin{aligned} \frac{\partial}{\partial \rho} \log q(\rho, \boldsymbol{V}) = & \sum_{n=1}^{N} \sum_{l=1}^{L} ( - \exp(\rho) B_{n,l} + \exp(\rho) \langle \log w_{n,l} \rangle \\ & - \psi(\exp(\rho)) \exp(\rho)) \end{aligned} \tag{17}$$

$$\frac{\partial}{\partial \boldsymbol{V}} \log q(\rho, \boldsymbol{V}) = -(\boldsymbol{E}_w \circ \exp(-\boldsymbol{X}\boldsymbol{V}))^\top \boldsymbol{X} + a_w \boldsymbol{X}^\top \boldsymbol{I}_{N,L} \tag{18}$$

where $\psi(x) = \frac{d}{dx} \log \Gamma(x)$, $\circ$ is the Hadamard product, $\boldsymbol{E}_w = (\langle w_{n,l} \rangle)$, and $\boldsymbol{I}_{N,L}$ is an $N \times L$ matrix of ones. $\log q(a_w, \boldsymbol{V})$ can be approximated by a normal distribution $N(\hat{\theta}, H^{-1}(\hat{\theta}))$, where $\hat{\theta}$ is the maximum a posteriori (MAP) estimator of $\theta$, and $H(\hat{\theta})$ is the Hessian of $\log q(a_w, \boldsymbol{V})$ evaluated at $\hat{\theta}$. For implementation, MAP estimates were obtained using the BFGS method in the optim function of R language. Credible intervals were computed using the optim function to compute the Hessian at the MAP estimates.

### 3. Handling missing data and choosing the number of communities

To handle missing data, we define a binary matrix $\boldsymbol{M} = (m_{n,k})$, where $m_{n,k} = 0$, if $y_{n,l}$ is missing, and 1 otherwise. The $p(\boldsymbol{S}|\boldsymbol{W}, \boldsymbol{V}, \boldsymbol{H})$ with missing data can be written as

$$
\begin{aligned}
p(\boldsymbol{S}|\boldsymbol{W}, \boldsymbol{H}) &= \prod_{n=1}^{N}\prod_{l=1}^{L}\prod_{k=1}^{K} p(s_{n,l,k}|w_{n,l}h_{l,k})^{m_{n,k}}. \\
&= \prod_{n=1}^{N}\prod_{l=1}^{L}\prod_{k=1}^{K} \exp(m_{n,k}(s_{n,l,k}\log(w_{n,l}h_{l,k}\tau_n) \\
&\quad - w_{n,l}h_{l,k}\tau_n - \log\Gamma(s_{n,l,k}+1))) \qquad (19)
\end{aligned}
$$

In this case, the update equations lead to the following:

$$
\begin{aligned}
q(s_{n,1:L,k}) \propto \exp\Big(\sum_{l=1}^{L}\Big(&s_{n,l,k}m_{n,k}(\langle\log w_{nl}\rangle + \langle\log h_{l,k}\rangle \\
&+ \log\tau_n) - \log\Gamma(s_{n,l,k}+1)\Big)\Big) \qquad (20)
\end{aligned}
$$

$$
\begin{aligned}
q(w_{n,l}) \propto \exp\Big(\Big(a_w + &\sum_{k=1}^{K} m_{n,k}\langle s_{n,l,k}\rangle - 1\Big)\log w_{n,l} \\
&- \Big(\langle B_{n,l}\rangle + \sum_{k=1}^{K} m_{n,k}\langle h_{l,k}\rangle\tau_n\Big)w_{n,l}\Big) \qquad (21)
\end{aligned}
$$

$$
q(\boldsymbol{h}_l) \propto \exp\left(\sum_{k=1}^{K}\Big(\alpha_k + \sum_{n=1}^{N} m_{n,k}\langle s_{n,l,k}\rangle - 1\Big)\log h_{l,k}\right), \qquad (22)
$$

where $q(s_{n,1:L,k})$ is the probability function of the multinomial distribution, $q(w_{n,l})$ is the density function of the gamma distribution, and $q(h_{l,k})$ is the density function of the Dirichlet distribution. The property of the gamma distribution and the Dirichlet distribution leads to the following:

$$
\langle w_{n,k}\rangle = \alpha_{n,k}^{(w)}/\beta_{n,k}^{(w)}, \qquad (23)
$$

$$
\langle\log w_{n,k}\rangle = \psi(\alpha_{n,k}^{(w)}) - \log\beta_{n,k}^{(w)}, \qquad (24)
$$

$$
\langle h_{l,k}\rangle = \frac{\alpha_{l,k}^{(h)}}{\sum_{k=1}^{K}\alpha_{l,k}^{(h)}}, \qquad (25)
$$

$$
\langle\log h_{l,k}\rangle = \psi(\alpha_{l,k}^{(h)}) - \psi\left(\sum_{k=1}^{K}\alpha_{l,k}^{(h)}\right), \qquad (26)
$$

where

$$\alpha^{(w)}_{n,k} = \langle a_w \rangle + \sum_{k=1}^{K} m_{n,k} \langle s_{n,l,k} \rangle \tag{27}$$

$$\beta^{(w)}_{n,k} = \langle B_{n,k} \rangle + \tau_n \sum_{k=1}^{K} m_{n,k} \langle h_{l,k} \rangle \tag{28}$$

$$\alpha^{(h)}_{l,k} = \alpha_k + \sum_{n=1}^{N} m_{n,k} \langle s_{n,l,k} \rangle \tag{29}$$

Using the delta method, $\langle a_w \rangle$ and $\langle \boldsymbol{B} \rangle$ can be approximated as follows [2]:

$$\langle a_w \rangle \approx \exp(\hat{\rho}) \tag{30}$$

$$\langle \boldsymbol{B} \rangle \approx \exp(-\boldsymbol{X}\hat{\boldsymbol{V}}). \tag{31}$$

To select the number of communities $L$, we replace some elements of $\boldsymbol{Y}$ with missing values, and we select a model by comparing the original value with the predicted value. The prediction accuracy was evaluated with the following test log-likelihood:

$$\text{test log-likelihood} = \sum_{(n,k)\in O} \log \text{Poisson}(y_{n,k} | \hat{y}_{n,k} \tau_n) \tag{32}$$

where $O$ is a set of element numbers of the missing test data, and $\hat{y}_{n,k}$ is the $(n, k)$-element of matrix $\boldsymbol{WH}$.

### 4. Implementation

We can avoid explicitly computing $\langle s_{n,l,k} \rangle$. Therefore, we can avoid storing the multidimensional array object. We define the following matrices:

$$\boldsymbol{L}_w = (\exp(\langle \log w_{n,l} \rangle)), \quad \boldsymbol{L}_h = (\exp(\langle \log h_{l,k} \rangle))$$

$$\boldsymbol{S}_w = \left( \sum_{k=1}^{K} \langle s_{n,l,k} \rangle \right), \quad \boldsymbol{S}_h = \left( \sum_{n=1}^{N} \langle s_{n,l,k} \rangle \right),$$

From Eq. (12), we can write

$$\sum_{k=1}^{K} \langle s_{n,l,k} \rangle = \sum_{k=1}^{K} y_{n,k} p_{n,l,k} \tag{33}$$

$$= \exp(\langle \log w_{nl} \rangle) \sum_{k=1}^{K} \frac{y_{n,k} \exp(\langle \log h_{l,k} \rangle)}{\sum_{l=1}^{L} \exp(\langle \log w_{n,l} \rangle) + \langle \log h_{l,k} \rangle)}. \tag{34}$$

Thus, we get

$$\boldsymbol{S}_w = \boldsymbol{L}_w \circ ((\boldsymbol{Y} \oslash (\boldsymbol{L}_w \boldsymbol{L}_h)) \boldsymbol{L}_h^\top). \tag{35}$$

Here, $\oslash$ denotes element-wise division. In the same manner as above, we obtain

$$\boldsymbol{S}_h = \boldsymbol{L}_h \circ (\boldsymbol{L}_w^\top (\boldsymbol{Y} \oslash (\boldsymbol{L}_w \boldsymbol{L}_h))). \tag{36}$$

The parameter-estimation procedure for BALSAMICO is summarized as follows:

(1) Set $\boldsymbol{\alpha}$
(2) Initialize $a_w$, $\boldsymbol{V}$, $\boldsymbol{E}_w = \boldsymbol{L}_w$, and $\boldsymbol{L}_h$.
(3) Iterate the following three steps until convergence:
   (a) Compute $\boldsymbol{S}_W$ and $\boldsymbol{S}_h$ from Eqs. (37) and (38), respectively.
   (b) Update $\boldsymbol{E}_w$, $\boldsymbol{L}_w$, and $\boldsymbol{L}_h$ from Eqs. (25), (26), and (28), respectively.
   (c) Update $a_w$ and $\boldsymbol{V}$ using Eq. (16).
